# AptViralDB: A Repository of Experimentally Validated Antiviral Aptamers

**DOI:** 10.64898/2026.07.08.737144

**Authors:** Nisha Bajiya, Shivansh Singh, Pushpendra Singh Gahlot, Gajendra P. S. Raghava

## Abstract

In an era of increasing drug resistance, exploring alternative molecules is crucial for the efficient management and treatment of viral diseases. Nucleic acid aptamers have emerged as highly promising candidates due to their exceptional target specificity, low immunogenicity, and versatile mechanisms for viral blocking. This manuscript describes AptViralDB, a manually curated database providing comprehensive information on experimentally validated antiviral aptamers. It contains 1,768 entries of antiviral aptamers against 40 viral species and 104 molecular targets, compiled from literature and existing databases. Each entry provides detailed annotations, including sequence, aptamer type, target, chemical modifications, binding affinity, antiviral activity, stability, and cytotoxicity. We also provide predicted secondary structures and their corresponding minimum free energy (MFE) values. Additionally, a knowledge graph created using ArcadeDB/openCypher enables users to seamlessly explore connections among aptamers, viruses, molecular targets, and biological activities. Finally, the platform offers advanced search and browsing tools, BLAST-based sequence similarity searches, GC-content analysis, downloadable datasets, and REST API access to support computational applications. (https://webs.iiitd.edu.in/raghava/aptviraldb/).

## 1. Introduction

Viruses remain among the most formidable infectious agents because of their rapid evolution, high mutation rates, and remarkable ability to evade host immune responses. Recurrent outbreaks caused by pathogens such as Severe acute respiratory syndrome coronavirus 2 (SARS-CoV-2), Human immunodeficiency virus (HIV), Influenza A virus, Zika virus, Ebola virus, and more recently emerging zoonotic viruses continue to pose major threats to global public health and socioeconomic stability (Bhadoria et al., 2021; Verikios, 2020; Keita et al., 2021). Although antiviral drugs and vaccines have substantially reduced disease burden, their long-term effectiveness is frequently compromised by viral genetic variability, antigenic drift and shift, drug resistance, pharmacokinetic limitations, adverse drug reactions, and drug-drug interactions(Duwe, 2017; Aw et al., 2025). In addition, many antiviral drugs are associated with adverse effects, including dermatological reactions, central nervous system disturbances, flu-like symptoms, blood cell-based irregularities, and organ toxicity, and may exhibit reduced efficacy due to drug-drug interactions (Morris, 1994; Akhvlediani et al., 2023; Zareifopoulos et al., 2020). These drawbacks emphasize the necessity for rapidly adjustable recognition systems capable of targeting similar viral elements with preserved selectivity.

Among emerging antiviral strategies, nucleic acid aptamers have emerged as promising candidates for antiviral therapeutics and diagnostics. Since the introduction of the Systematic Evolution of Ligands by EXponential enrichment (SELEX) technology in 1990 (Robertson and Joyce, 1990; Ellington and Szostak, 1990; Tuerk and Gold, 1990), a variety of aptamers targeted to purified viral proteins, intact virions, virus-infected cells, and even host proteins of the virus were developed (Ilgu and Nilsen-Hamilton, 2016). Depending on the intended application, specialized SELEX strategies, including Protein-SELEX, have enabled the isolation of aptamers recognizing viral envelope glycoproteins, spike proteins, reverse transcriptases, RNA-dependent RNA polymerases, proteases, nucleocapsid proteins (Quang et al., 2017). Whole-virus SELEX preserve native conformational epitopes, while Cell-SELEX enables recognition of virus-infected cells. The integration of next-generation sequencing technologies into SELEX workflows has enabled rapid aptamer discovery, as able to analyze the functionality of sequenced aptamers and identify the active sites of the aptamers with antiviral properties using enriched aptamer libraries (Takahashi, 2018).

Unlike conventional antiviral molecules, aptamers can interfere at different levels in the viral life cycle. They can block viral attachment by hindering receptor recognition, inhibit membrane fusion with the cell and viral entry, suppress genome replication by targeting viral polymerases or reverse transcriptases, interrupt viral protein maturation, recognize the virusinfected host cells, and serve as molecular probes for highly sensitive diagnostics and biosensing platforms (Zou et al., 2019; Kim and Lee, 2021; Krüger et al., 2021; Chakraborty et al., 2022). Their ease of chemical synthesis, low immunogenicity, high reproducibility, and specific chemical modification further expand their utility for targeted drug delivery and therapeutic development (Ni et al., 2021; Chen et al., 2021). The various mechanisms through which antiviral aptamers interfere with viral infection and replication are outlined in Figure 1. This opens a vast array of possibilities for aptamers and their use in the development of next-generation anti-infective agents.

**Figure 1:**
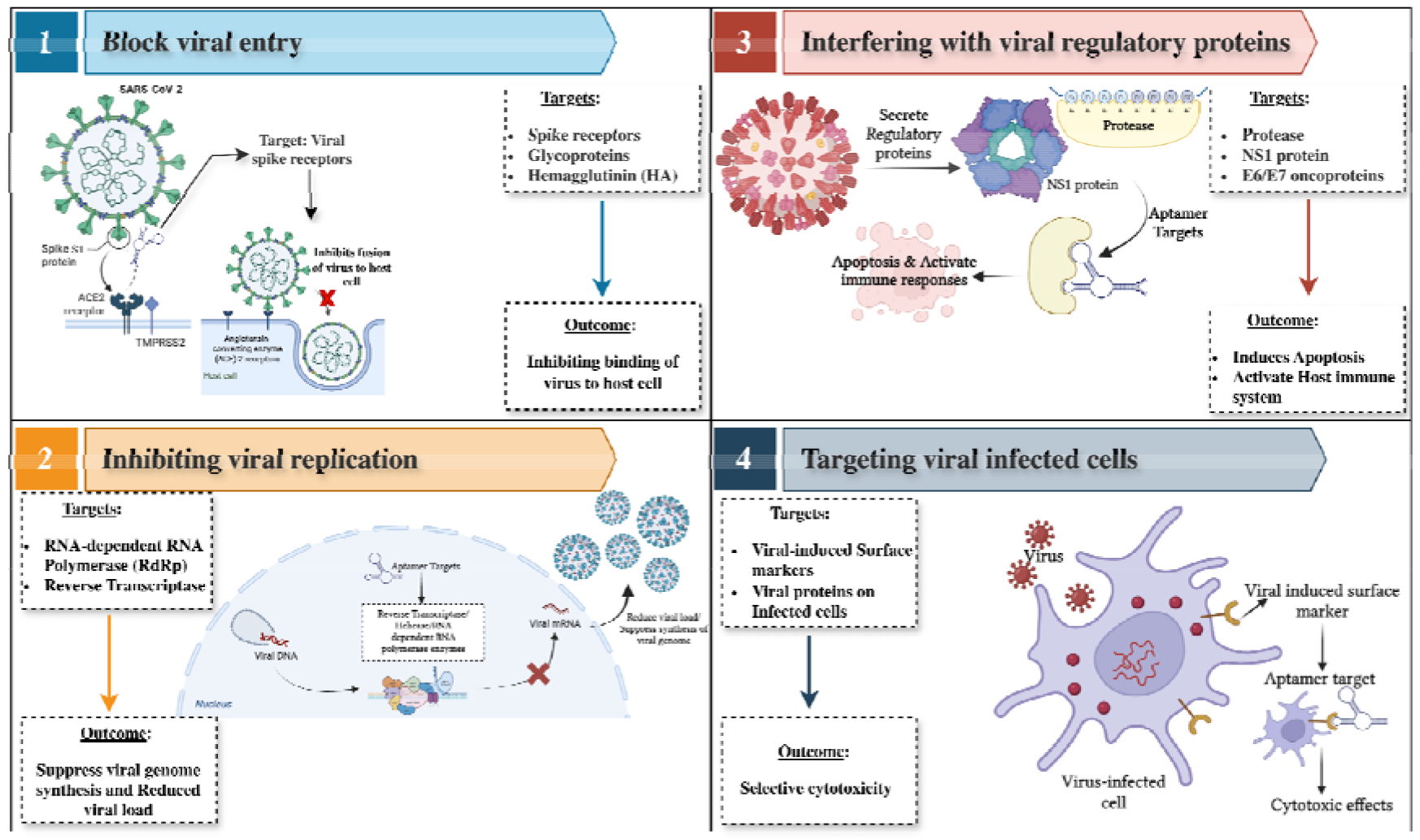
Diagrammatic representation of the antiviral mechanism by antiviral aptamers.

Antiviral aptamers display excellent target specificity due to the folding of aptamers into intricate 2-D structures (e.g. Stem-loops, hairpins, bulges) and 3-D structures (e.g. Pseudoknots, G-quadruplexes) that result in significant conformational changes and unique conformational rearrangements complementary to their targets or the viral structure upon binding (Micura and Höbartner, 2020). Structural elucidation using X-ray crystallography, NMR spectroscopy, and computational modeling has revealed that aptamer-target interactions are governed by molecular shape complementarity along with various noncovalent interactions such as hydrogen bonding, van der Waals forces, hydrophobic effects, and electrostatic forces (Zhang et al., 2021). Following selection, techniques such as Surface Plasmon Resonance (SPR), Bio-layer Interferometry (BLI), Microscale Thermophoresis (MST), fluorescence-based assays, and enzyme-linked binding assays is commonly performed to quantify binding affinity to estimate dissociation constants (Kd) which are in pico- to nanomolar range comparable to those of antibody (Hasegawa et al., 2016), inhibitory concentration values (IC50), and effective concentration values (EC50) (Pawel et al., 2022; Malysheva et al., 2024). Then, the antiviral efficacy of these identified aptamers is usually evaluated using viral neutralization assays, plaque reduction assays, antiviral cell-based assays, and experiments on inhibition of viral entry.

With increased emphasis on antiviral aptamers research, a variety of computational methods such as the prediction of aptamer secondary and tertiary structure, molecular docking, molecular dynamics simulation, and artificial intelligence (AI) based approach have been introduced for more efficient discovery and optimization of aptamers, which predictaptamer folding, aptamer-target interaction and the conserved motif of the aptamers before experimental validation (Yu et al., 2019; Lee et al., 2023; Fang et al., 2024; Ruiz-Ciancio et al., 2024; Malysheva et al., 2024). However, the success of these methods relies heavily on high-quality, well-curated, experimentally validated antiviral aptamers.

The necessity of well-annotated datasets has already been demonstrated in the field of antiviral peptides, where dedicated databases and prediction tools (Qureshi et al., 2014; Gupta et al., 2016; Patiyal et al., 2020; Zhang et al., 2022; Ullah et al., 2024; Duy and Srisongkram, 2025; Sun et al., 2025; Nawaz et al., 2026) have accelerated sequence analysis, activity prediction, and the design of new antiviral peptides. However, computational tools dedicated to antiviral aptamers are still limited. Although several general-purpose aptamer databases have been developed (Lee et al., 2004; Cruz-Toledo et al., 2012; Askari et al., 2024; Chen et al., 2024; Lu et al., 2026), they are not exclusively focused on antiviral applications and generally lack comprehensive viral classification, standardized binding data, and detailed functional annotations. Therefore, comprehensive comparisons, data-driven discovery, and the development of robust AI-based antiviral aptamer predictive algorithms remain challenging.

To date, only four clinical trials involving antiviral or virus-associated aptamers have been registered on ClinicalTrials.gov, reflecting the clinical translational gap. These include two trials for COVID-19, one for the treatment evaluation using ApTOLL (NCT05293236), a DNA aptamer targeting TLR4 and a saliva-based DNA aptamer diagnostic platform for its detection (NCT04974203), third, a tenofovir aptamer-based biosensor used to monitor adherence to HIV drug (NCT04870671), and fourth, BC 007 (NCT05911009), a 15-mer ssDNA aptamer drug which is being evaluated as a therapy for long COVID. This limited progression can be attributed to challenges such as nuclease instability, delivery barriers for intracellular viral targets, pharmacokinetic optimization, and evolving regulatory pathways for nucleic acid therapeutics. Moreover, relevant antiviral aptamer data are dispersed across heterogeneous studies, with inconsistent data reporting, which hampers the systematic comparison across viral species and the development of robust computational prediction tools, underscoring the need for a centralized and structured database.

To bridge this gap, we developed AptViralDB, a comprehensive repository of experimentally validated antiviral aptamers that provides a structured, searchable platform for the global scientific community. The platform provides advanced search and browsing, downloadable datasets, knowledge graph (KG)-based relationship exploration, REST API access, and integrated analysis tools, thereby establishing a computational resource that supports largescale bioinformatics analyses, machine learning model development, and the rational discovery of next-generation antiviral aptamers.

## 2. Materials and Methods

### 2.1. Data Collection

To curate antiviral aptamer entries, we conducted PubMed and Google searches using keywords such as “Antiviral aptamer”, “viral aptamer”, “therapeutic aptamer”, “diagnostic aptamer”, etc. Also, we have retrieved data from existing databases, including the UTexas Aptamer Database (Askari et al., 2024), AptaBase (https://www.iitg.ac.in/proj/aptabase/about.html), and AptaDB (Chen et al., 2024). Each retrieved article and review was manually screened to ensure relevance. After removing duplicates and unnecessary articles and keeping experimentally validated antiviral aptamers, the search returned articles from 1992 to 2025, totalling 364 PMIDs and 17 DOIs. When multiple variants of an aptamer were reported, each variant was reported as a separate entry. The selected publications were then thoroughly reviewed for the collection of antiviral aptamer data utlized as antivirals, diagnostics, and targeted delivery. The complete repository framework is shown in Figure 2.

**Figure 2:**
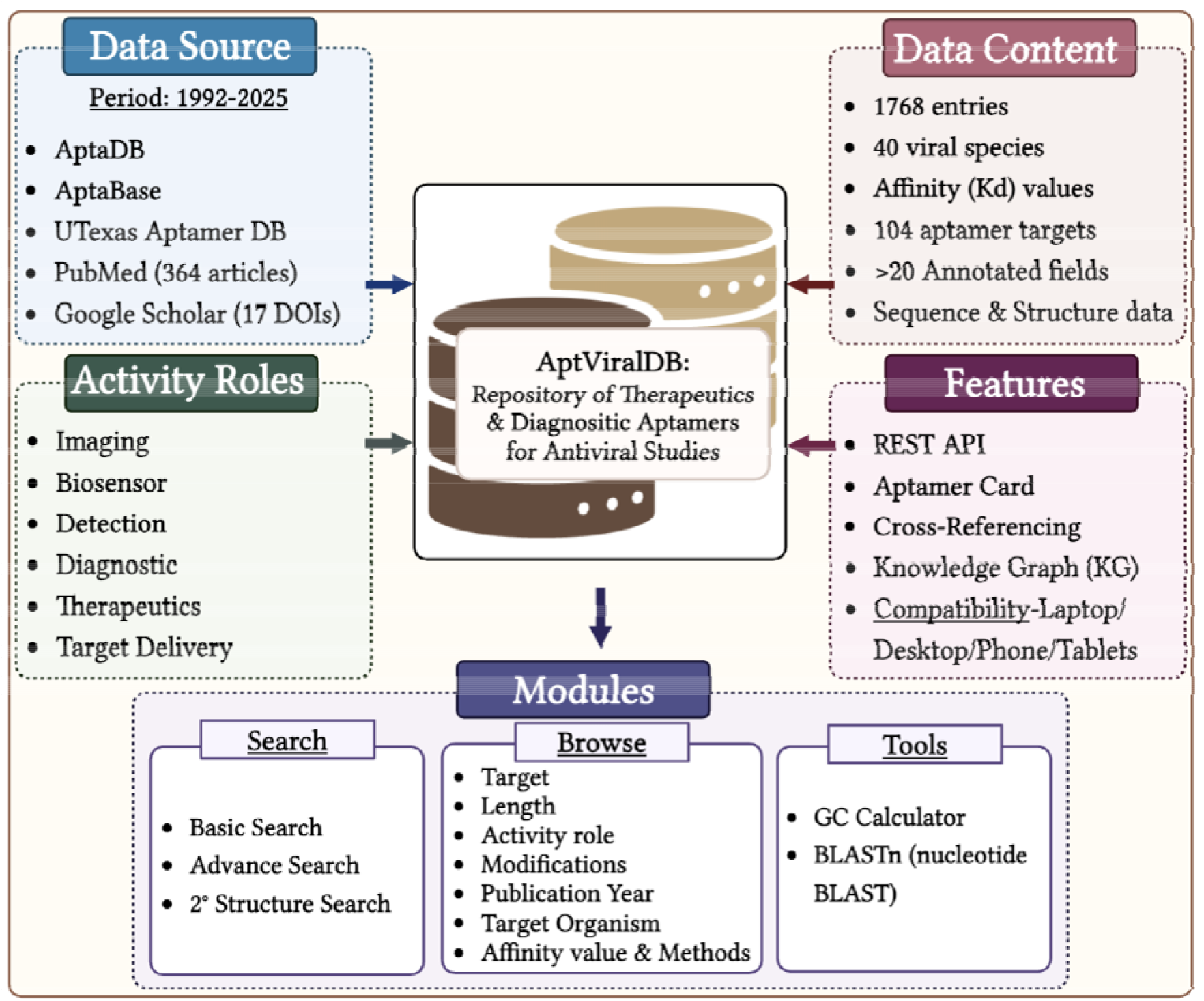
The Architecture of AptViralDB.

### 2.2. Data Annotation and Standardization

For each entry, detailed information about the aptamer was manually extracted, including its sequence, length, type, chemical modifications, target name, target virus, experimental SELEX method, library used and number of rounds performed to obtain the desired aptamer, affinity determination method, and binding affinity (Kd). Additional information, such as the study’s objective, the inhibition outcome (IC50 or EC50), diagnostic applications, and the activity role (therapeutic, detection, etc.), was recorded. In addition, cytotoxicity, stability, half-life, and patent information were curated when available. All reported sequences are in uppercase, and modified nucleotides were consistently annotated along with the type of modification. The repository comprises 40 different virus classes and 104 reported aptamer targets. The entries have been linked to their corresponding aptamer cards, which contain detailed, curated information. Also, optimal secondary structures of the aptamers were predicted using the ViennaRNA Package 2.0 (Lorenz et al., 2011) with minimum free energy (MFE) in kcal/mol, and their representation in standard Dot-notation and structural elements, including stems, loops, hairpins, etc., was reported.

### 2.3. Database Architecture and Implementation

AptViralDB was developed using a Relational Database Management System (RDBMS) architecture implemented in MySQL (version 5.5.62) and Apache HTTP Server (version 2.4.7). The backend was built with PHP, while the frontend was designed with TypeScript, HTML5, React and Tailwind CSS to ensure compatibility across modern web browsers. The database framework was organized into interconnected tables storing aptamer details, virus type, target information, experimental methods, and publication metadata for querying, browsing, and analyzing data. Users can access it for free at https://webs.iiitd.edu.in/raghava/aptviraldb.

### 2.4. Web Interface and Functional Modules

The web interface was designed to provide automatic access to data. Users can perform a simple search using virus name, target name, or aptamer ID. An advanced search module allows filtering based on multiple parameters, including aptamer type, target virus, aptamer targets, and affinity range. The database also provides browsing options categorized by ‘Target organism-wise’, ‘Target-wise’, ‘Length-wise’, ‘Modification-wise’,’ Affinity-wise’, ‘Activity role’, ‘Affinity Determination method’, and ‘Publication year’. Additionally, Similarity search using nucleotide BLAST and the GC calculator tool have been incorporated into the analysis. To obtain more information about the aptamer incorporated from other repositories, such as AptaDB and Aptabase, users can access our “Cross-Referencing” module. For programmatically downloading the data, the “REST API” module has also been provided. Additionally, the user can download the sequence data in FASTA format and the predicted secondary structures in SVG format to facilitate further analysis.

### 2.5. Knowledge Graph Construction and Integration

To enable a structured representation of complex relationships among antiviral aptamers and their related data, we adopted a knowledge graph (KG) based on the ArcadeDB multi-model graph database (https://arcadedb.com/knowledge-graphs.html) within AptViralDB. During data preprocessing, curated records were presented as tables, mapped to structured subject-predicate-object triples (head-relation-tail), and stored as CSV files. Three core triple sets were constructed: (1) Aptamer-TARGET-Viral Protein (capturing aptamers and direct interactions with the viral protein targets), (2) Aptamer-TARGETS_ORGANISM-Virus (connecting aptamers to the viral organisms being targeted), and (3) Aptamer-HAS_ACTIVITY_ROLE-Functional Role (capturing the purpose of the aptamer). The CSVs were then loaded into the ArcadeDB instance through OpenCypher LOAD CSV queries utilizing MERGE operations so as to avoid node deduplication across files, defining 4 vertex types (Aptamer, Target, Organism, ActivityRole) connected via 3 directed edge types. This architecture enables multi-level traversal between relationships, thereby enabling exploration of novel findings such as viral families sharing the same target, evaluating the effects of modifications in specific roles, and assessing the repurposing potential of existing aptamers in other viruses. The ArcadeDB platform has enabled the construction of an interoperable data structure that supports advanced analytical workflow via its Studio interface and API, as well as a flexible architecture for integrating further data and analyses, including predictive modeling and machine learning.

## 3. Results and Discussion

### 3.1 Data Analysis

AptViralDB currently contains 1768 experimentally validated antiviral aptamers targeting 40 different viral species, with 121 entries present in UTexas databases, 106 in AptaDB and 26 in Aptabase. The exponential growth in cumulative entries from 1992 to 2025 is depicted in Figure 3. After 2011, a significant increase in the number of published studies can be observed, which may reflect increased scientific attention to the antiviral applications of aptamers, triggered by major global outbreaks such as the 2009 influenza pandemic and the 2021 COVID-19 pandemic, which stimulated rapid development of novel molecular recognition systems as a potential basis for diagnostics or therapeutics (Bhadoria et al., 2021). Various oligonucleotide types have been used to construct antiviral aptamers (Table 1) and can be categorized into ssDNA, RNA and xeno nucleic acid (XNA) based aptamers. The majority (69.1%) are DNA aptamers, as reflected in higher stability compared to RNA aptamers due to enhanced resistance to hydrolysis and easier handling and use (Ni et al., 2021). There is a limited proportion (2.0%) of XNA-based aptamers (i.e. FANA, HNA, TNA aptamers), which are a group of chemically modified nucleic acid analogs, created with the aim to increase resistance towards nuclease and improve stability toward antiviral applications while maintaining high affinity binding to their targets (Rose et al., 2019).

**Table 1:** The distribution of aptamers based on the type of nucleic acids in AptViralDB.

| Aptamer Type | Count | Percentage (%) |
| --- | --- | --- |
| ssDNA | 1221 | 69.1 |
| ssRNA | 510 | 28.86 |
| ssXNA | 36 | 2.04 |

**Figure 3:**
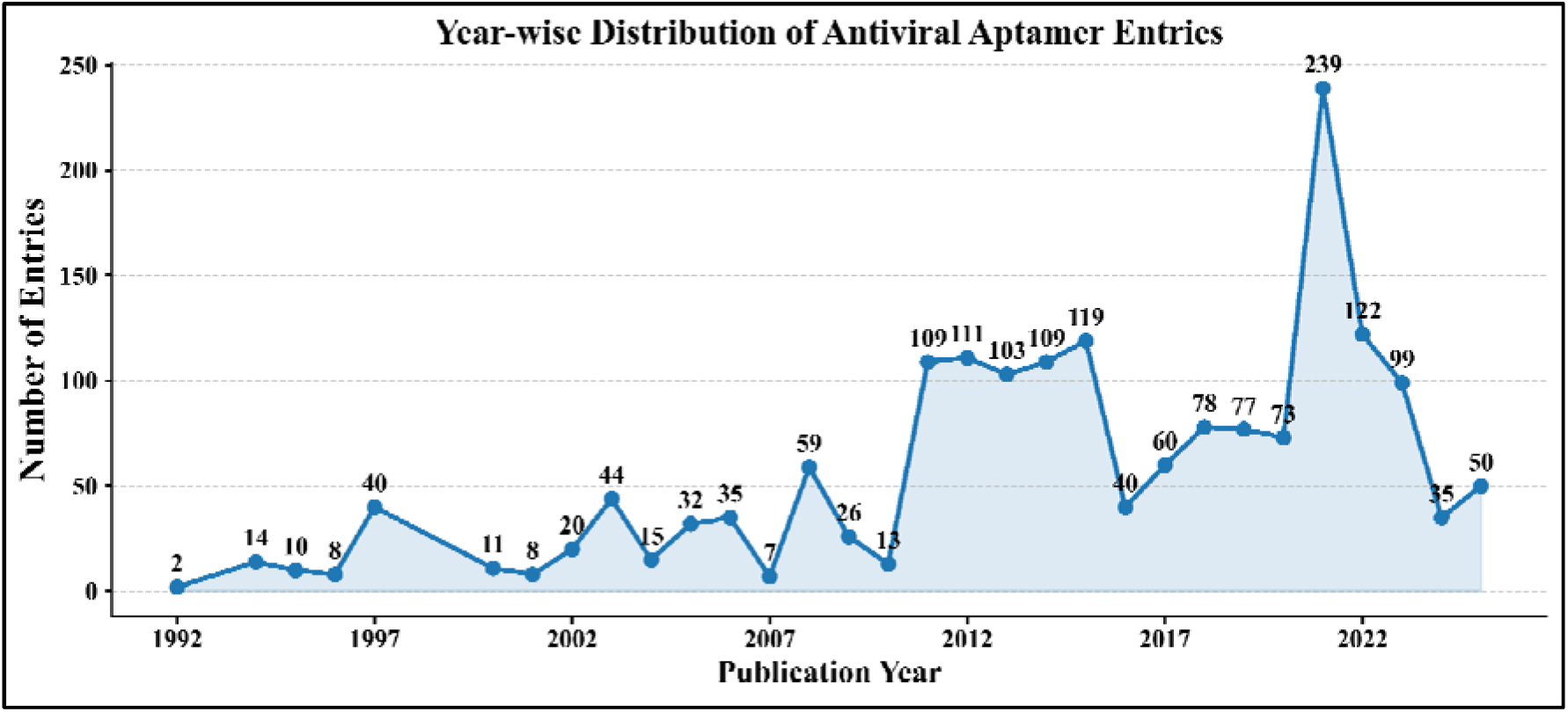
Line plot showing the cumulative year-wise growth trend of aptamers in antiviral studies.

Figure 4 (A) demonstrates that analyzing the target organism reveals that coronaviruses, HIV, influenza, and hepatitis viruses account for the majority of entries, aligning with global disease burden and historical antiviral research priorities (Duwe, 2017; Aw et al., 2025). Similarly, in Figure 4 (B), most aptamers in the database target whole viral cells with unspecific target information (12.21%) and surface glycoproteins such as hemagglutinin, spike proteins, and receptor-binding domains, as well as intracellular enzymes like reverse transcriptase, reflecting the established strategy of blocking viral attachment, entry and replication machinery (Zou et al., 2019; Chakraborty et al., 2022).

**Figure 4:**
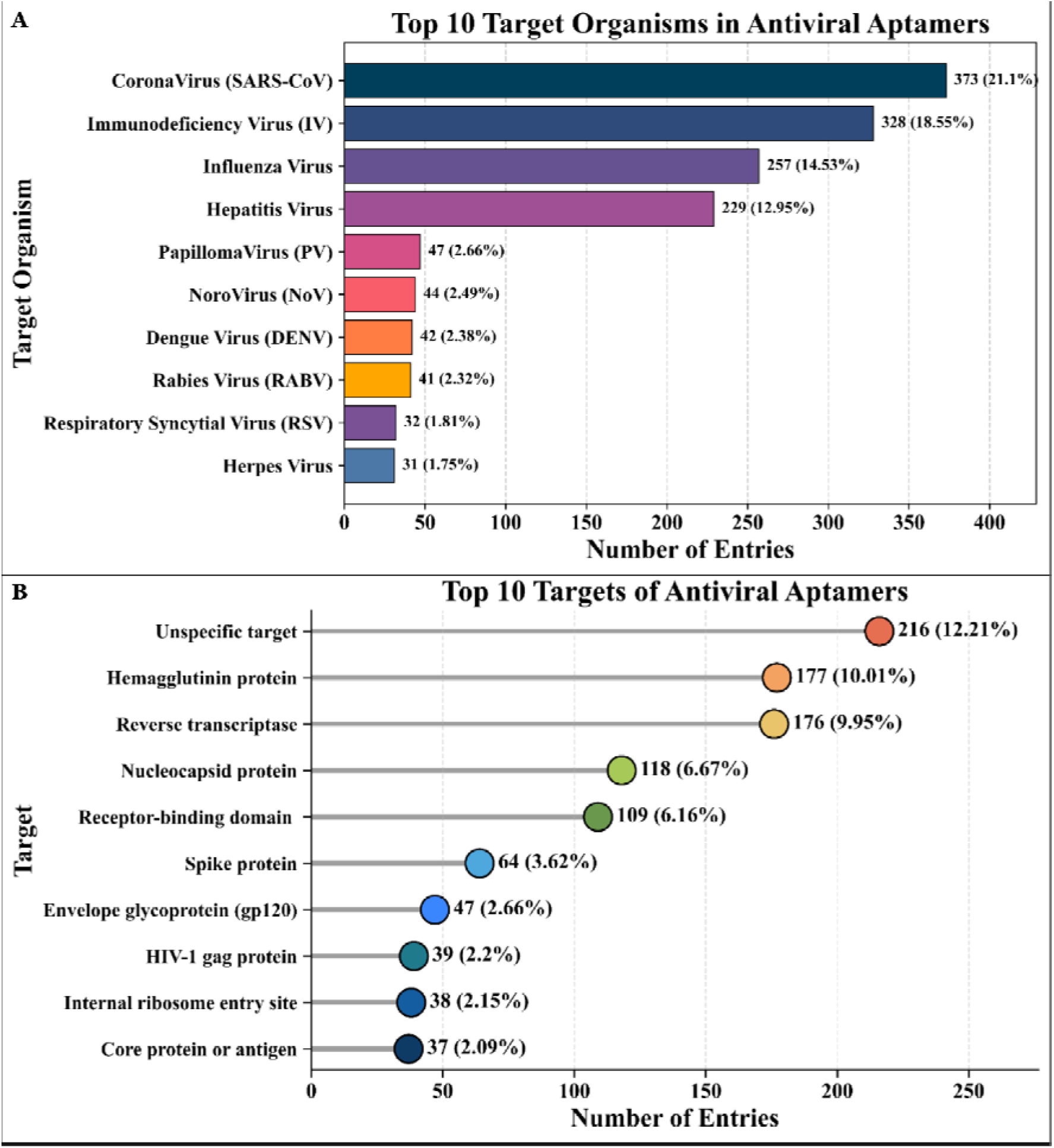
Bar plots showing the (A) Top 10 virus species reported in AptViralDB, (B) Top 10 most commonly used viral targets for aptamer studies reported in AptViralDB.

Length-wise distribution indicates that 80% of antiviral aptamers fall within the 41-100 nucleotide range, particularly for therapeutic and biosensor applications (Figure 5 (A)), which is commonly reported as optimal for maintaining structural complexity and high-affinity binding while preserving synthesis efficiency (Hasegawa et al., 2016; Zhang et al., 2021). Regarding affinity validation from Figure 5 (B), Surface Plasmon Resonance is the most frequently used method, consistent with its recognition as a gold-standard, real-time, label-free technique for kinetic characterization of aptamer-target interactions (Pawel et al., 2022).

**Figure 5:**
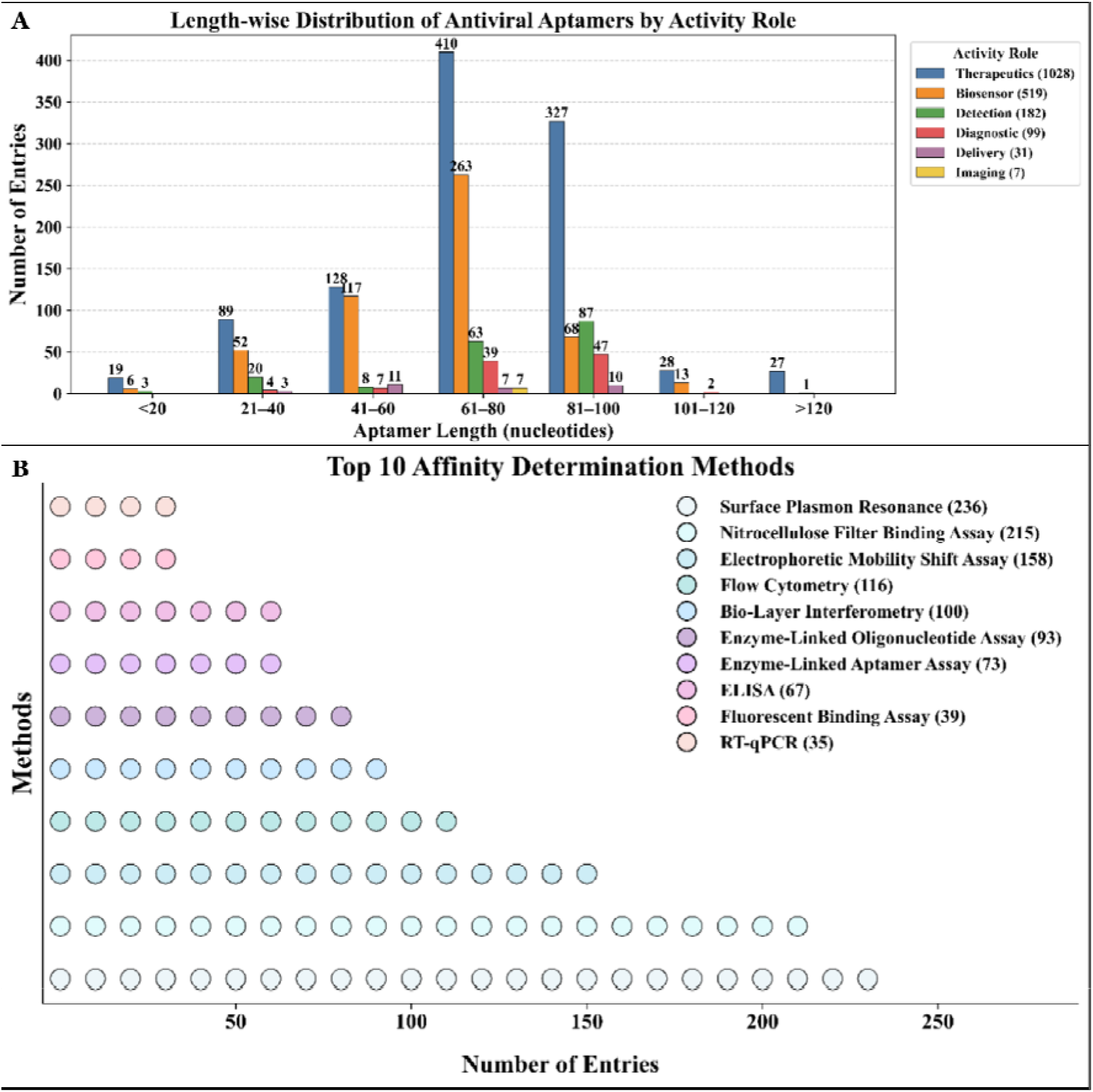
(A) Bar plot showing the distribution of activity roles by aptamer length. (B) The top 10 most commonly used methods for affinity determination.

#### Thermodynamic Stability and Secondary Structure Analysis of Antiviral Aptamers

Aptamer secondary structure has been well characterized by experimental methods, including X-ray crystallography, cryo-EM, NMR, SAXS, and CD spectroscopy, which provide structural and dynamic details but need advanced instrumentation and laborious protocols (Malysheva et al., 2024). Whereas secondary structure prediction using computational tools is a quick and scalable method for determining base-pairing patterns and folding stability, which will guide subsequent experimental validation (Sinharay et al., 2026). As depicted in Figure 6 (A), the highly negative correlation coefficient between sequence length and MFE (R = -0.579) shows that a longer sequence of aptamers possesses more thermodynamically stable secondary structures (i.e., more negative MFE), since the length of nucleotides offers potential for greater base pairing and complex folding in both RNA and DNA molecules (Yang et al., 2025). The unimodal distribution of MFE with an average value of -10.44 kcal/mol (Figure 6 (B)) indicates most aptamers have moderate secondary structure stability, while a few aptamers possess highly negative MFE values, which signifies the stronger folding stability. Besides, it has been shown that ssRNA aptamers generally have lower MFE values than ssDNA aptamers because of greater structural variability and intramolecular interactions, as recently supported by experimental studies on comparative energetic stabilities of nucleic acid folding (Pal and Levy, 2019). Furthermore, evaluation of dot-bracket notation showed a considerably large percentage of paired bases, which implies the presence of relatively compact secondary structures and stem-loops as the most predominant motifs.

**Figure 6:**
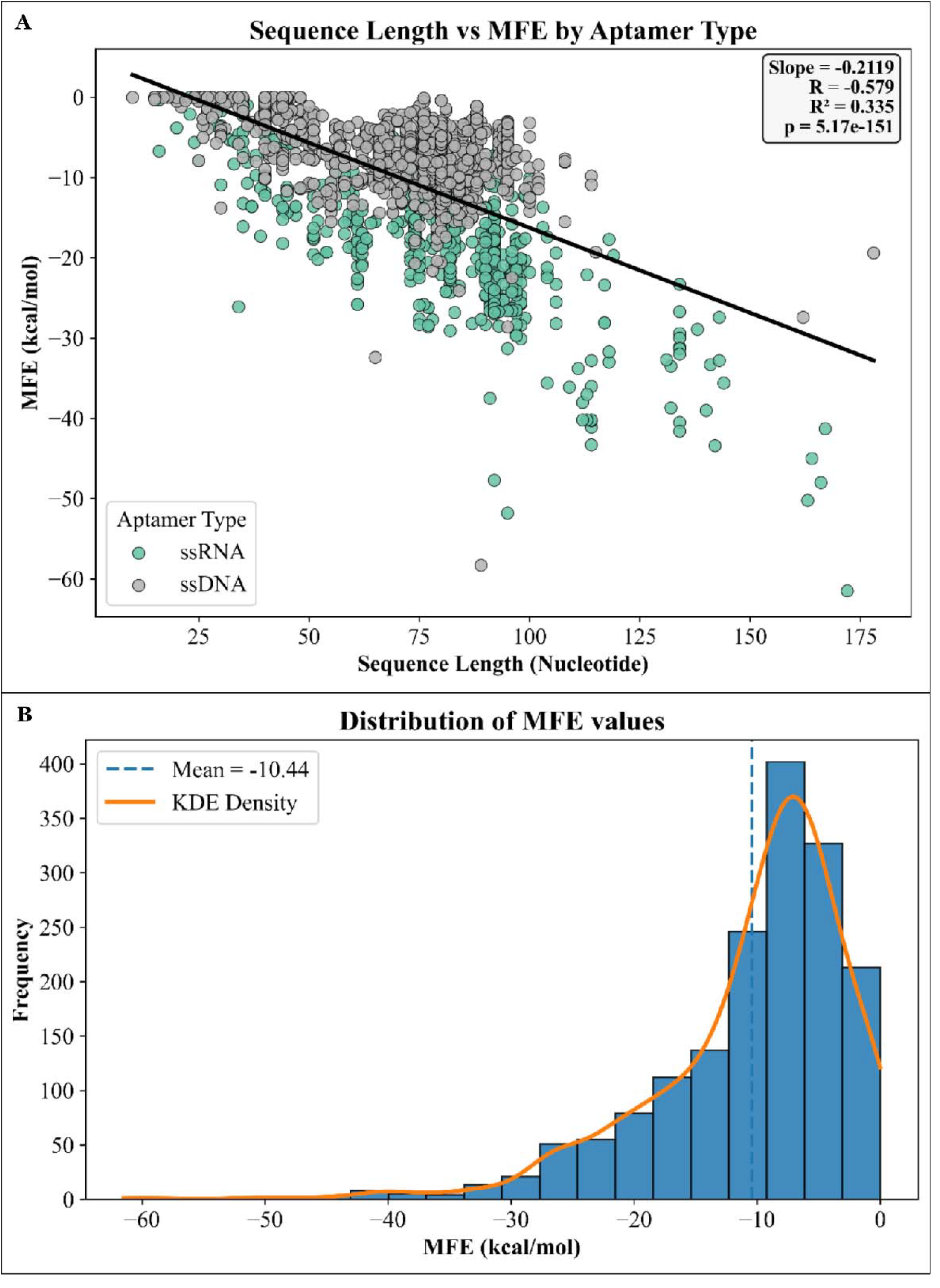
(A) Regression plot between aptamer sequence length and predicted MFE values. (B) Histogram plot showing the distribution of predicted MFE values in the dataset.

Overall, these observations suggest that the aptamers in the dataset possess stable, structured configurations that could facilitate efficient, high-affinity binding to their viral targets.

## 4. Conclusion

In brief, AptViralDB is a comprehensive, manually curated database of experimentally validated antiviral aptamers. By aggregating scattered literature into a structured, searchable platform, this resource addresses a current gap in antiviral research. It provides a valuable resource for researchers engaged in antiviral drug discovery, diagnostic development, and computational modeling. The curated dataset can be used for structure-function analysis, motif identification, binding affinity prediction, and the development of machine-learning-based methods to identify new antiviral aptamers. Furthermore, the repository can support rapid response efforts during emerging viral outbreaks by enabling aptamer repurposing and comparative analysis.

## Author Contributions

NB collected, curated and analyzed the data. SS developed the web server’s front-end and back-end. The manuscript was prepared by NB, SS, and GPSR. The idea for the project was conceived and coordinated by GPSR. The final version of the paper was reviewed and approved by all the authors.

## Acknowledgements

The authors are thankful for the necessary facilities and infrastructure, as well as fellowships and financial support, provided by the Council of Scientific and Industrial Research (CSIR), the University Grants Commission (UGC), the Department of Biotechnology (DBT), and the Department of Computational Biology at IIITD, New Delhi. We also like to thank Draw.io for making the Figures.

## Data availability

We will update it soon.

